# Accuracy of phylogenetic reconstructions from continuous characters analyzed under parsimony and its parametric correlates

**DOI:** 10.1101/2024.01.03.574081

**Authors:** Victor A Tagliacollo, Mario de Pinna, Junior Chuctaya, Alessio Datovo

**Author notes:** Corresponding author, VAT.

## Abstract

Quantitative traits are a source of evolutionary information often difficult to handle in cladistics. Tools exist to analyze this kind of data without subjective discretization, avoiding biases in the delimitation of categorical states. Nonetheless, the ability of continuous characters to accurately infer relationships is incompletely understood, particularly under parsimony analysis. This study evaluates the accuracy of phylogenetic reconstructions from simulated matrices of continuous characters evolving under alternative evolutionary processes and analyzed by parsimony. We generated 100 trees to simulate 9,000 matrices containing 26 terminals and 100 continuous characters evolving under: Brownian-Motion (BM), Ornstein-Uhlenbeck (OU) and Early-Burst (EB) processes assuming variable parametrizations. Our comparisons of cladograms revealed that matrices analyzed by parsimony carry phylogenetic signals to infer relationships, but the accuracy is higher for matrices simulated under BM, regardless of the parameterization schemes. Implementation of equal or implied weighting with multiple penalization strengths against homoplasies did not affect cladogram inferences. Accuracy of continuous characters in resolving relationships is skewed toward apical nodes of the trees. Our simulations provide controlled tests of the usefulness of quantitative traits in phylogenetics, specifically under neutral evolution, and demonstrate their effectiveness in estimating shallower nodes among recently diverged species, regardless of parameters and weighting schemes.

## BACKGROUND

Cladistics is an evolutionary approach usually associated with the application of parsimony to infer relationships among species by analyzing homologous characters and identifying derived shared states to establish monophyletic groups (J. S. Farris, 1983; J. S. Farris et al., 1970; Hennig, 1966). It requires that characters are ontologically independent entities and variation is discretized into mutually exclusive states (J. S. Farris, 1983; J. S. Farris et al., 1970; Sereno, 2007). Nevertheless, inappropriate discretization or subjective thresholds can introduce analytical biases that affect tree accuracy, often leading to erroneous conclusions about species relationships (Goloboff et al., 2006; Thiele, 1993). To avoid arbitrary choices, alternative methods have been proposed to convert continuous variation into discrete character states (Kitching et al., 1998; Archie, 1985; Goldman, 1988; Thiele, 1993; Wiens, 2001), although their effectiveness and limitations are debated (J. S. Farris, 1990; Goloboff et al., 2006).

Most phenotypic characters are heritable features that provide empirical evidence of shared ancestry among species over time. The aim of phylogenetic analysis is to discriminate the level(s) at which such shared similarities are homologous (i.e., synapomorphies). Characters can be classified as either qualitative (e.g., qualitative or categorical morphological characters, nucleotides) or quantitative (e.g., morphometrics, meristics)(Thiele, 1993). While qualitative characters are prevalent in cladistics, thresholds between phenotypic character states may be subject to debate, including bias in character selection and subjectivity in state delimitations (Wiens, 2001; Zelditch et al., 2000). Although often downplayed in cladistic studies (e.g., MacLeod, 2002), quantitative traits also provide important insights into the evolutionary history of species (Klingenberg & Gidaszewski, 2010; Rae, 1998; Thiele, 1993; Wiens, 2001; Zelditch et al., 2000). Some phylogenetic analysis programs allow direct incorporation of quantitative traits, quantified along a finite continuous spectrum, as a data source for tree inference (Goloboff & Catalano, 2016; Höhna et al., 2016). These programs objectively delimit character states for tree searches (Catalano et al., 2010; Goloboff et al., 2006; Goloboff & Catalano, 2011), thus avoiding potential biases associated with subjective choices of discretization thresholds between states (J. S. Farris, 1990). Nevertheless, incorporating quantitative traits into phylogenetics requires the assumption of an underlying evolutionary process of character change (e.g., neutral evolution) that is not explicitly stated in parsimony reconstructions. Therefore, simulation-based experiments assuming explicit evolutionary models are useful to provide insight into the effectiveness of continuous traits in accurately inferring species relationships analyzed by parsimony (e.g., Huelsenbeck & Hillis, 1993, but see Goloboff et al., 2018).

Explicit models can be used to infer evolutionary processes and simulate character changes over time. For quantitative traits, these evolutionary models are more commonly characterized by Gaussian distributions and describe memoryless stochastic movements of phenotypic changes in continuous time and morphospace (Blomberg et al., 2020). Among the most widely used stochastic models for simulating character evolution are Brownian motion and its derivatives, including Ornstein-Uhlenbeck and Early-Burst (e.g., Parins-Fukuchi, 2018b, 2018a; Wright, 2019). These models simulate a variety of evolutionary scenarios of character change, including random drift, selection, and phenotypic variations expected in events of adaptive radiations (Álvarez-Carretero et al., 2019). These models have been widely used and have been applied to the study of macroevolutionary patterns to understand the underlying processes of character evolution in different groups of organisms (Álvarez-Carretero et al., 2019). By generating simulated data matrices of continuous characters, those models allow investigating the impact of quantitative traits on cladistic analyses, providing insights into the dynamics of evolutionary change. Testing the evolutionary process of data matrices may help to understand how accurate phylogenetic inferences are on the basis of continuous traits (Parins-Fukuchi, 2018b).

This study uses simulations to investigate the accuracy of cladogram reconstruction by parsimony when analyzing continuous characters not arbitrarily discretized and under alternative weighting schemes. By simulating character matrices under different evolutionary models, we provide evidence for the utility of continuous traits in inferring relationships at varying node depths, regardless of the rate of character evolution and the regimens of homoplasy down-weighting. Through comparisons of cladogram reconstruction, we circumscribe conditions under which quantitative characters are a valuable source of evolutionary signal for phylogenetic studies.

## METHODOLOGY

### Simulations

This study used the R package phytools (Revell, 2012) to generate 100 stochastic trees under a pure-birth model, with a speciation (birth) rate of 1.0 and a relative time of 1.0 for consistency across all trait simulations. Each tree comprised 25 terminals plus one outgroup bound without edge length, which served as a standard rooting point. We used these trees to simulate matrices of 100 continuous characters evolving stochastically along their edge lengths under three time-continuous processes: unbounded Brownian-Motion (BM) (Beaulieu et al., 2012; Butler & King, 2004; Hansen, 1997), Ornstein-Uhlenbeck (OR) (Felsenstein, 1973, 1985; Gingerich, 1993) and Early-Burst (EB) (Harmon et al., 2003, 2010). All character simulations started with an ancestral state at the root set to 0.0. Table 1 summarizes the parameter values in the simulations of potential biological scenarios of continuous character evolution.

**Table 1.** Parameter inputs to simulate matrices of continuous characters under three time-continuous processes of character state evolution. All character simulations started with an ancestral root state of 0.0. Sigma2 (σ2) = evolutionary rate. Omega (θ) = optimal value. Alpha (α) = Adaptive rate. Beta (β) = decay rate.

|  |  |  |  |  |  |
| --- | --- | --- | --- | --- | --- |
|  | **Brownian-Motion (BM)** |  |  |  |  |
| $\sigma^2$ | 0.05 | 0.50 | 1.00 | 1.50 | 3.00 |
|  | **Ornstein-Uhlenbeck (OU)** |  |  |  |  |
| $\sigma^2$ | 0.50 | | | | |
| $\theta$ | 0.00 | | | | |
| $\alpha$ | 0.05 | 0.50 | 1.00 | 1.50 | 3.00 |
|  | **Early Burst (EB)** |  |  |  |  |
| $\sigma^2$ | 0.50 | | | | |
| $\beta$ | -0.05 | -0.50 | -1.00 | -1.50 | -3.00 |

Under the BM model, we generated matrices of characters evolving through unidirectional random drifting, with variance increasing unboundedly over time, controlled by alternative sigma-squared (σ2) rates (Table 1). This process describes the evolution of neutral changes over time among characters not under selection. For the OU model, we generated matrices of characters evolving under selection towards an optimal adaptive zone. This process was controlled over time by alternative alpha (α) rates, which specified pull strengths towards an optimal θ value (Table 1). Lower α rates implied lower variance and greater homoplasy. Under EB, we generated matrices emulating scenarios of character evolution expected in events of adaptive radiations (*sensu* Simpson, 1953). This Gaussian, time-varying process was controlled by multiple decay beta (β) rates slowing variance over time (Table 1). Larger absolute β values implied faster decay in character changes over time. For the OU and EB models, we applied a σ2 rate of 0.5 across all character matrices because rate values had no significant impact on the overall accuracy of cladogram reconstructions (see Results). To prevent unbalanced dominance of characters in higher orders of magnitude, we rescaled values between 0.0 and 1.0 (Koch et al., 2015). An overview of the simulation steps is shown in Figure 1. Datasets are available as supplementary material.

**Figure 1.**
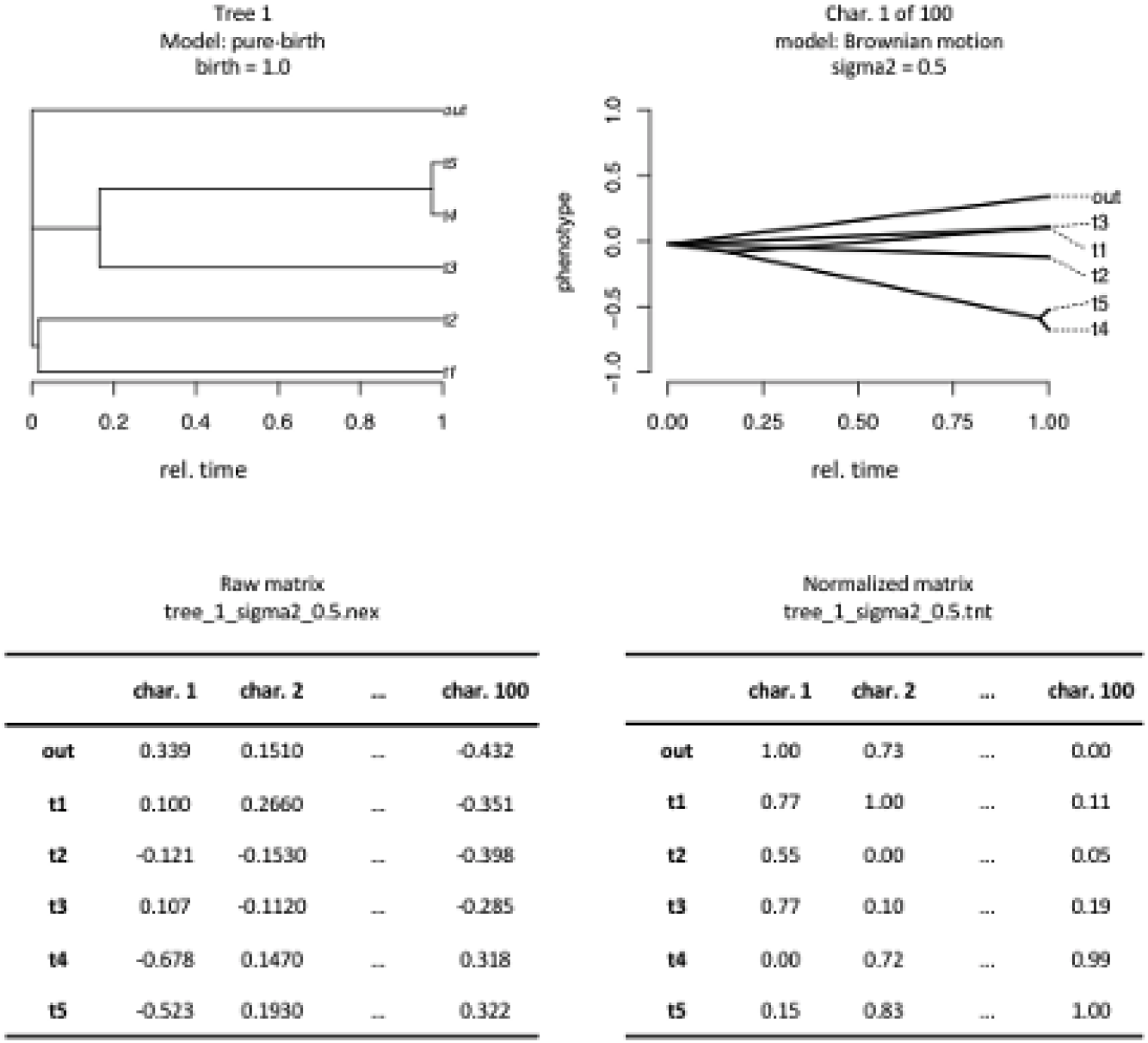
Outline of the simulation steps used to evaluate the accuracy of phylogenetic reconstructions from continuous characters analyzed by maximum parsimony in TNT. A) Simulation of a pure-birth tree assuming a speciation (birth) rate of 1.0, relative time scale of 1.0, and including an outgroup without edge length and five more terminals. B) Simulation of one continuous character evolving along edges of a pure-birth tree under an unbounded Brownian Motion (BM) time-continuous process with an evolutionary sigma2 (σ2) rate of 0.5 and ancestral root state set to 0.0. C) Schematic character matrix showing one hundred simulations of continuous traits under BM model. D) Rescaled character matrix of continuous characters with value ranges between 0 and 1.

In our simulations, we treated continuous characters as evolving under constant-rate conditions, with the same rate parameter applied uniformly across edges resulting in the accumulation of variance between sister lineages since their last common ancestor. Our simulations treated characters as independent entities with no covariance in any dimensionality, in accordance with the phylogenetic assumption of character independence (Sereno, 2007). In empirical matrices, continuous characters can exhibit covariance, especially at low dimensionality with a few coevolving characters (Adams & Felice, 2014). However, phylogenetic inferences from simulated matrices of continuous characters have shown that topological accuracy from correlated traits are as accurate as those from uncorrelated traits, except at very high dimensionality and covariance (Parins-Fukuchi, 2018b). Additionally, covariance between continuous characters can be eliminated by PC transformation, thus avoiding potential bias due to large covariant modules (see e.g., in (González-José et al., 2008); but criticism in (Adams et al., 2011).

### Phylogenetic inferences in TNT

For cladogram reconstruction by parsimony, we analyzed a batch of files containing rescaled matrices in TNT v. 1.5 (Goloboff & Catalano, 2016). Given the number of terminal taxa (26), searches for the most parsimonious trees (MPTs) were performed using the heuristic RAS+TBR strategy, with 500 replicates, holding 10 trees per replication, and random seed set to zero. After this first round of searches, the MPTs retained in memory were submitted to a second round of TBR searches to ensure that all MPTs (up to the limit of 10,000 trees) of the sampled tree islands were found (thus eliminating possible tree ‘overflows’ of the initial searches). We analyzed the matrices using both equal weighting (EW) and implicit weighting (IW) searches. EW parsimony searches for the trees with shortest length imposing the same weight to all characters, regardless of their degree of homoplasy. IW parsimony applies differential weighting against homoplasy by searching for the trees with maximum Fit (Goloboff, 1993). The strength of character down-weighting is controlled by the concavity constant k (Goloboff, 2014). The lower the value of k, the stronger its down-weighting of homoplasy. We tested six k values – 0.5, 1, 2, 3, 5, and 10 – to cover a reasonable range of down-weighting regimes of homoplastic characters.

### Accuracy of cladogram reconstructions

We performed comparisons of cladograms by computing topological errors and node congruences between simulated pure-birth trees and cladogram reconstructions from the simulated matrices. Using Robinson-Foulds (RF) symmetric distances (Robinson and Foulds 1981), we reported topological errors by counting shared and unshared bipartitions between rooted trees, excluding branch length estimates. RF distances are presented in relative scale, with lower values closer to 0.0 indicating higher similarity between compared trees. We illustrated the RF values using boxplots created in the ggplot2 package (Wickham & Chang, 2016)

We computed node congruences among cladograms over relative time by mapping the results onto pure-birth trees using two strategies. One strategy applied majority-rule consensus to summarize cladogram reconstructions from character matrices generated under single evolutionary processes with multiple rate parameters. We represented the frequencies of node congruences as percentages, with values closer to 100.0 indicating higher overlap among compared nodes. We illustrated node congruences using scatterplots with smoothed contour density estimates generated in the ggplot2 package (Wickham & Chang, 2016). In a second strategy, we applied an index of node similarity using element-based scores, an optimized version of the Jaccard index implemented in the ViPhy program (Bremm et al., 2011). The scores are presented on a relative scale, with values closer to 1.0 indicating higher similarity between compared clades.

## RESULTS

### Cladogram reconstruction under alternative models and parametrizations

RF distances indicate that matrices of continuous characters do carry phylogenetic signals for inferring species relationships, but the accuracy of node reconstructions is dependent on the evolutionary time-continuous processes used to simulate character matrices (Fig. 2). Overall, reconstruction errors are lower for character matrices generated under BM compared to those using OU or EB processes, regardless of the parameterization in the simulations (Fig. 2). RF distances also indicated that continuous characters performed consistently across multiple rate parameters, independently of the chosen character evolution models (Fig. 2).

**Figure 2.**
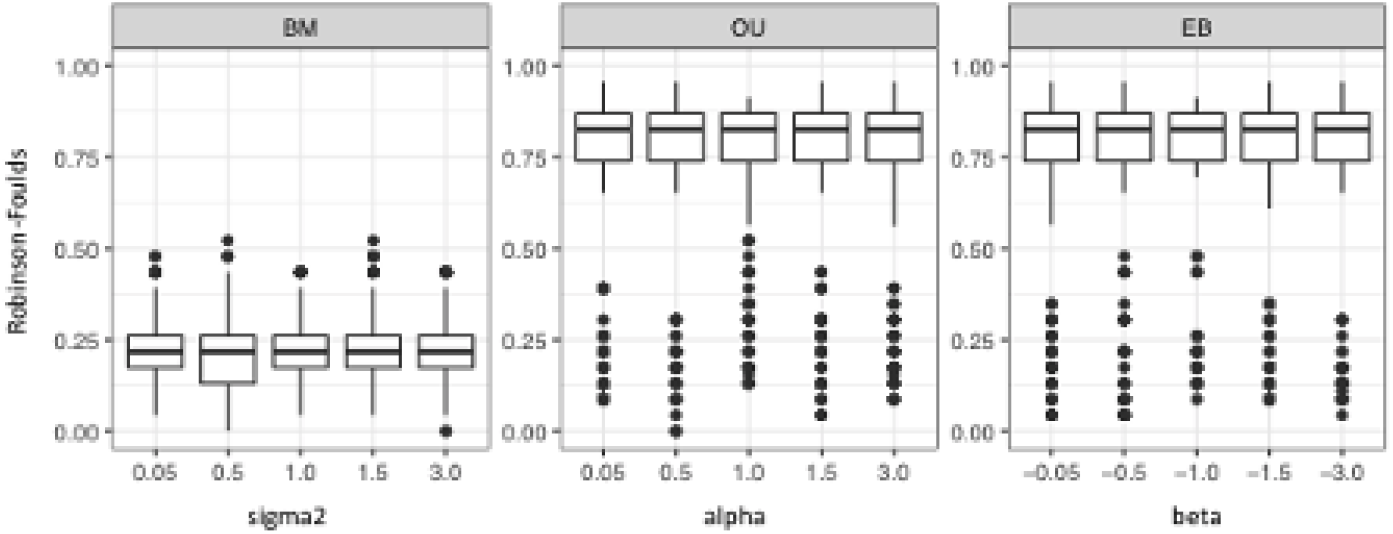
Accuracy of cladogram inferences from matrices simulated under three time-continuous models of character evolution and analyzed by maximum parsimony in TNT. Topological errors calculated by Robinson-Foulds (RF) symmetric distances suggest that continuous characters carry informative evolutionary signals to cladogenetic relationships. The accuracy of cladogram reconstruction is significantly higher from matrices simulated under Brownian Motion (BM) when compared to Ornstein-Uhlenbeck (OR) and Early-Burst (EB) models, regardless of the evolutionary (sigma2), adaptive (alpha), or decay (beta) rate parameters, respectively. RF values closer to 0.0 indicate higher similarity between compared cladograms.

### Cladogram inferences under alternative weighting schemes

RF distances indicated that neither EW nor IW schemes, the latter regardless of k values, affected the accuracy of cladogram reconstructions across simulations using any evolutionary model (e.g., BM, OU, or EB) and parametrizations (e.g., sigma2, alpha, and beta) (Fig. 3). In other words, the differential weighing of homoplasies derived from continuous characters does not interfere with the accuracy of cladogram reconstructions by parsimony. In all cases, the accuracy of inferences from character matrices generated under BM was higher than those generated by OU or EB processes (Fig. 2 and 3).

**Figure 3.**
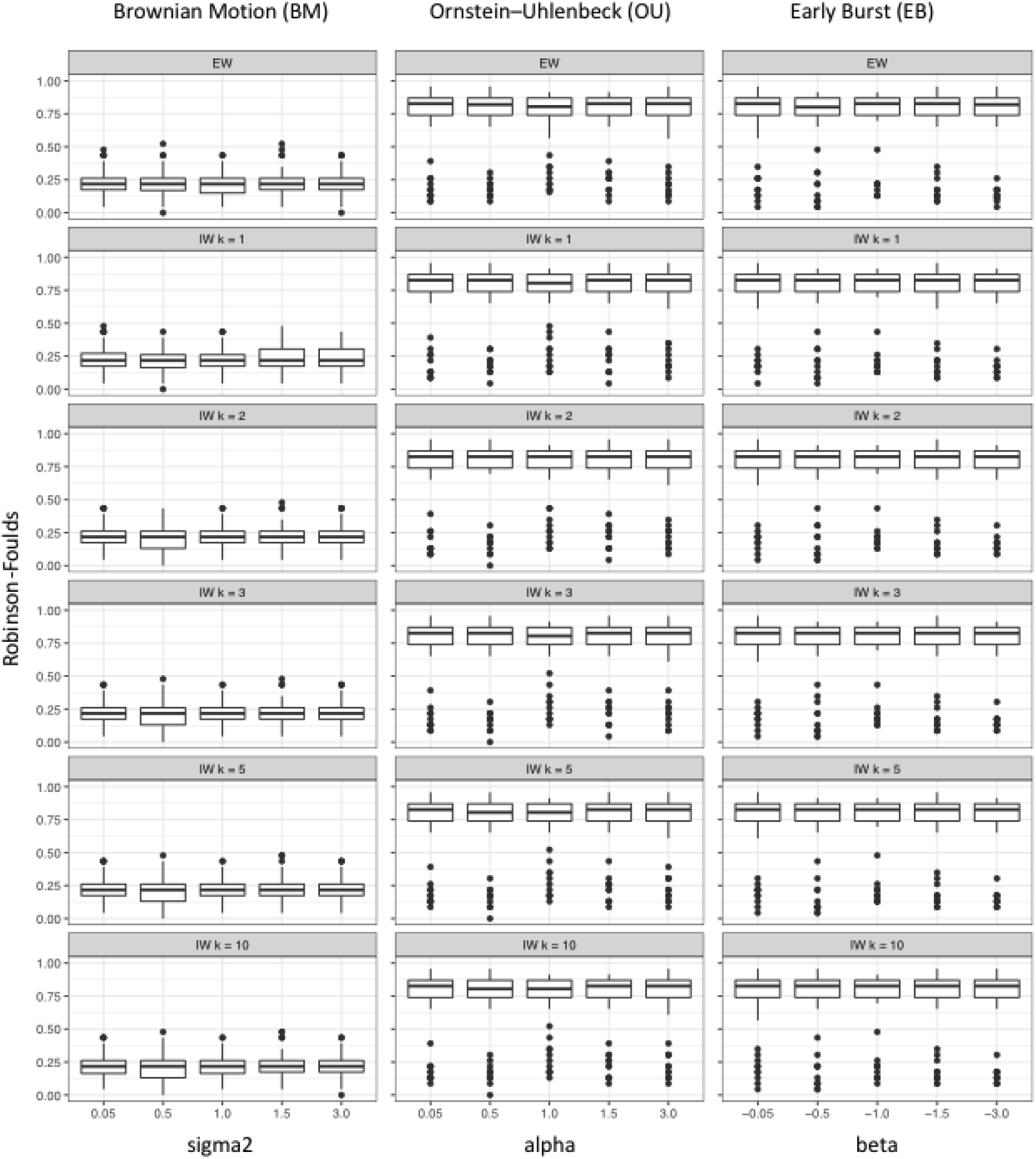
Alternative weighting strategies and accuracy of cladogram inferences from matrices simulated under three time-continuous models of character evolution and analyzed by maximum parsimony in TNT. Topological errors calculated by Robinson-Foulds (RF) symmetric distances suggest that neither Equal Weighting (EW) nor Implied Weighting (IW) schemes, the latter with alternative k values, interfere in accuracy of cladogram inferences across simulations, regardless of the evolutionary models (i.e., BM, OU, or EB) and their respective rate parameters (i.e., sigma2, alpha, and beta). RF values closer to 1.0 indicate higher dissimilarity between compared cladograms.

### Node congruence across relative tree length

Node congruence indicated that cladogram reconstructions using characters evolving under the BM process more accurately recovered relationships at both intermediate and shallow node heights, whereas inferences using characters evolving under the OU and EB processes were accurate only at shallow nodes (Fig. 4). These results indicate that continuous characters analyzed by parsimony can more accurately infer relationships at shallower nodes, particularly between recently diverged sister species (Fig. 5), regardless of whether characters evolve under random drift (i.e., BM), selection (i.e., OU) or adaptive radiation (i.e., EB). However, at intermediate nodes close to 0.5 of the tree height, only BM can more often estimate accurate relationships (Fig. 4 e 5). Our results suggest that continuous characters analyzed by parsimony do not perform well in inferring relationships at deeper nodes closer to the root (Fig. 4 e 5).

**Figure 4.**
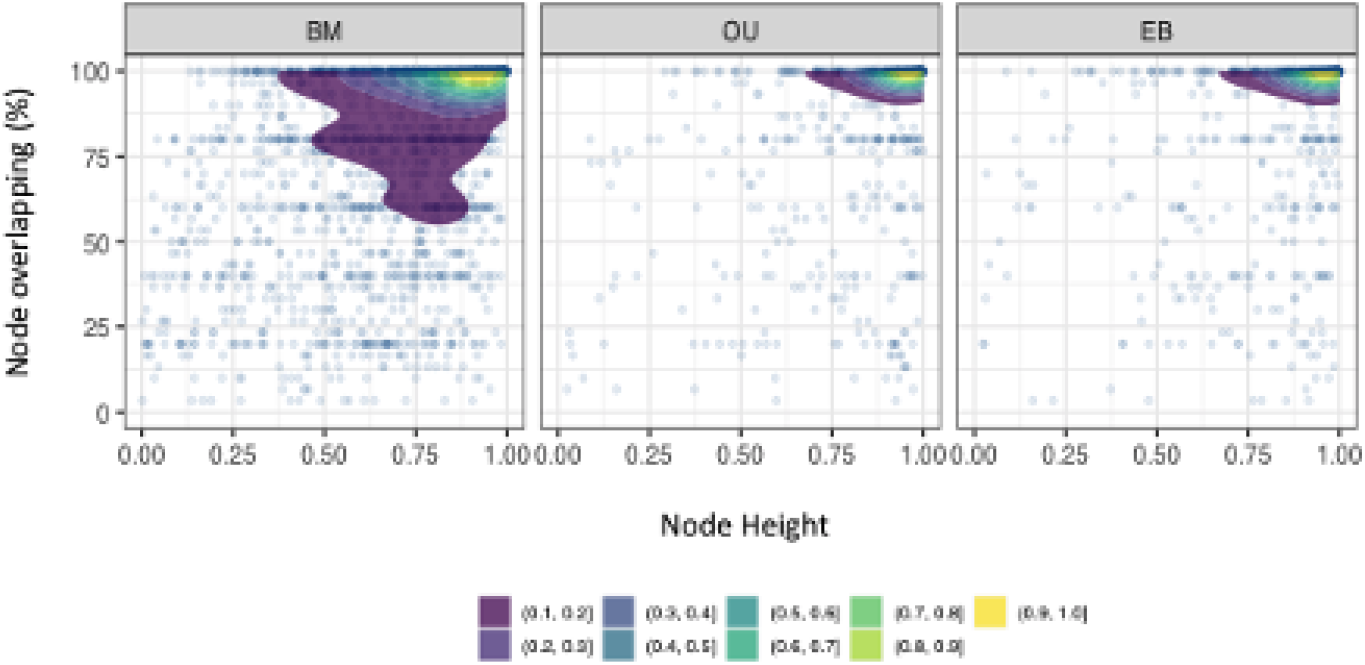
Node congruence of cladogram reconstructions summarized by majority-rule consensus across relative time scale under three evolutionary processes with multiple rate parameters. Overall, simulations of continuous characters using the BM process recover a larger number of correct relationships and at intermediate node height compared to the OU and EB processes. These results suggest that continuous characters are sources of phylogenetic information to resolve species relationships at shallower nodes, with higher accuracy when characters are evolving under random drifting, rather than selection (OU) or during adaptive radiations (EB).

**Figure 5.**
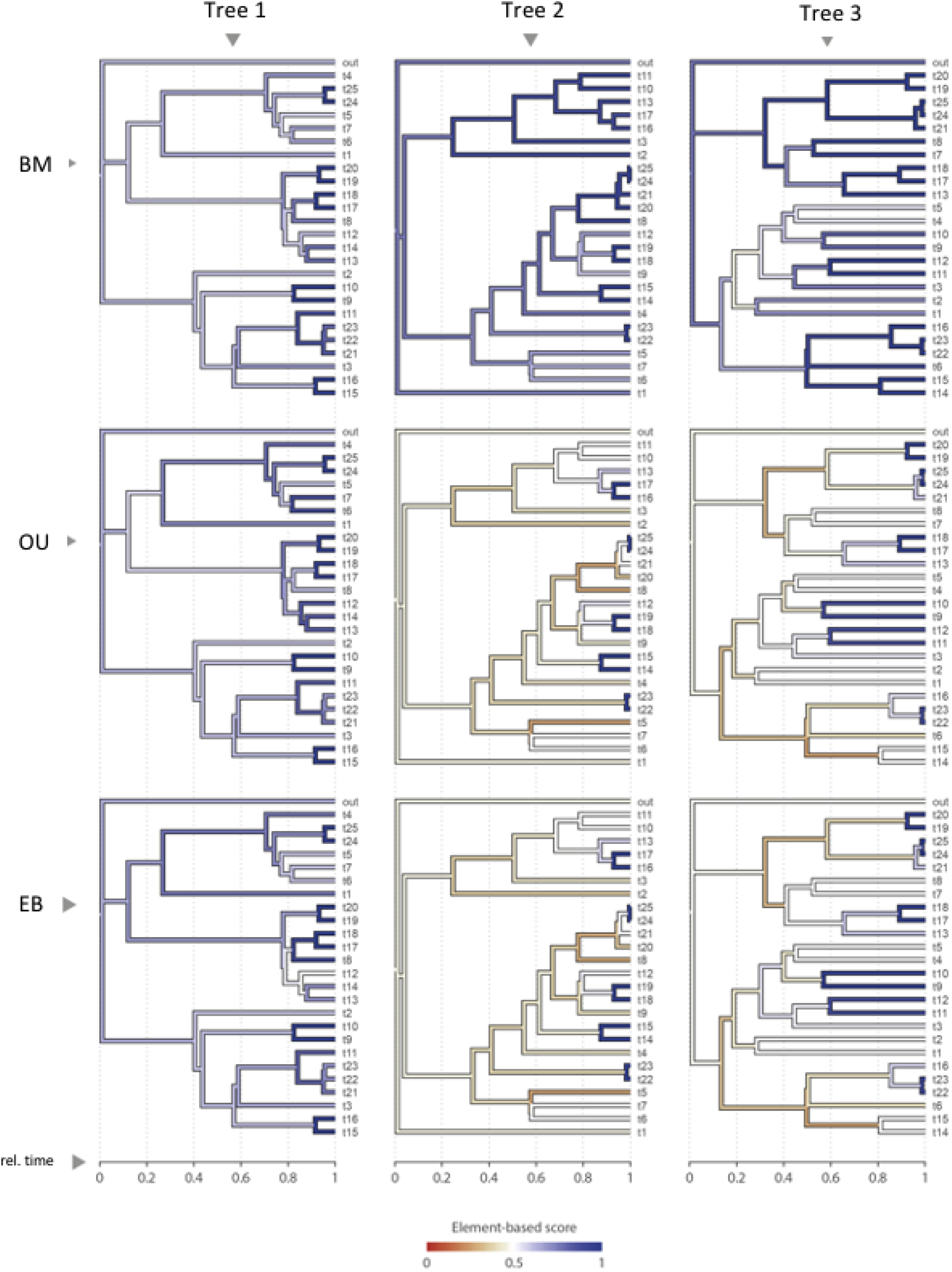
Effectiveness of continuous characters in the inference of shallower nodes of cladograms. Continuous characters carry phylogenetic signals and more often resolve relationships at shallower nodes, particularly between sister taxa splitting more recently in relative time. Element-based scores with darker blue gradient color indicate higher clade similarity among compared trees.

## DISCUSSION

### Cladogram reconstruction under alternative models and evolutionary rates

The accuracy of cladogram reconstruction from continuous characters depends on the evolutionary process applied to simulate the character matrices. Our study indicates that parsimony reconstruction errors are often lower for characters evolving under random drift (Brownian motion) compared to the other evolutionary processes (e.g., Ornstein-Uhlenbeck and Early burst). In our simulations, characters generated under this neutral evolutionary process were more accurately recovered by the parsimony algorithm, regardless of parametrization and weighting schemes. Characters simulated by directional selection or accelerated evolution yielded quantitatively similar outcomes, resulting in weaker phylogenetic signals that sometimes hinder rather than improve tree reconstructions by parsimony, especially at deeper node levels.

Our results also show that the observed topological reconstructions are often statistically indistinguishable regardless of the evolutionary rates. This implies that the rate of evolution has minimal influence on parsimony-based cladogram reconstructions, whereas the underlying process of character evolution is more critical. The potential cause for the efficiency of parsimony in recovering diverse evolutionary rates may be attributed to the strategy of rescaling the continuous characters on a relative scale (Goloboff et al., 2006; Goloboff & Catalano, 2016), which results in the homogenization of their amplitudes, thereby minimizing topological differences due to rate changes (Koch et al., 2015).

### Cladogram inferences under alternative weighting schemes

Our findings further suggest that the accuracy of phylogenetic reconstructions using continuous traits under parsimony is essentially unaffected by alternative weighting schemes against non-uniform homoplasies equal or implied weighting – even in the tests with relatively strong down-weighting strengths that typically produce distortions in empirical studies (i.e., k = 0.5 and 1). There are at least two possible explanations for this, which are not mutually exclusive: either quantitative traits display low levels of evolutionary convergence or reversal, or discretization without subjectivity results in descriptively minimal homoplasy, making it difficult to detect convergent evolution. This minimal homoplasy makes it challenging to detect and down-weight those exhibiting convergent evolution. The indifference of implied weighting (IW) with continuous characters is also demonstrated with empirical matrices employing an alternative tree distance index (Koch et al., 2015). Even though topological differences may be discernible, statistical outcomes from both EW and IW are equally probable, and, in such cases, only additional characters can contribute to a convergence towards more robust relationships.

Convergence of continuous characters is unlikely due to their stochastic nature, which means that they rarely take values that can be interpreted as convergent in parsimony-based cladograms. A criticism of the use of continuous random variables in cladistics relates to the biological significance of probabilistic distribution means (Pimentcl & Riggins, 1987; Rae, 1998). The challenge is to discretize continuous characters into meaningful states without hindering our understanding of convergence patterns. While character states evolving under selection often converge to optimal peaks, the discretization schemes used by the programs may not treat such peaks as a single convergent state. For continuous characters, convergence may not always be detected because states with values that are close but numerically distinct are not treated as convergent and thus are not penalized by the weighting schemes. Therefore, continuous characters undergoing selection or adaptive radiation events may introduce noise into phylogenetic analyses, which may ultimately affect the inference of relationships between distantly related species.

The indifference of alternative weighting regimes in parsimony analyses of qualitative data contrasts with studies that have evaluated only categorical data. These analyses show that implied-weighted parsimony often outperforms equal weighting (Goloboff et al., 2008, 2018). It is worth noting that our simulations did not evaluate the effects of combining different evolutionary processes and rates. Nevertheless, it seems reasonable to assume that the accuracy of parsimony-based cladogram reconstructions will be lower when multiple processes are combined, compared to those generated from data matrices under the random drifting processes.

### Node congruence across relative tree length

Our simulations indicate that cladogram reconstructions with continuous characters evolving under random drifting exhibit greater accuracy for intermediate and shallow node heights (Figs. 4, 5). Conversely, inferences using continuous characters evolving under selection or adaptive radiation demonstrate accuracy primarily for shallow nodes between closely related sister species (Figs. 4, 5). Overall, morphological characters often exhibit faster data saturation and often result in reduced phylogenetic signals (Dávalos et al., 2014). Characters evolving under random drift are more efficient in reconstructing deeper nodes, although their resolution is seldom greater than about half the relative time of the cladograms. This observation indicates that continuous characters are especially valuable for determining relationships among recently diverged groups or when the evolutionary process is driven by random drift, resulting in resolutions that usually do not exceed about half of the topology. Relying solely on continuous characters in cladistics may lead to a low-resolution topology, especially for nodes close to the tree root. To overcome this limitation, it is crucial to integrate data from diverse sources into the analysis, with each source providing information that may help resolve different regions of the cladogram (e.g., (Calegari et al., 2019; Mirande, 2019; Pastana et al., 2022; Presti et al., 2023; Terán et al., 2020).

Even at slow evolutionary rates, the accuracy of inferences using continuous data is limited to specific parts of the tree, rarely achieving high accuracy for the entire tree. In the case of selection and adaptive radiation models, resolution is even more limited to apical nodes, especially among sister groups. Therefore, caution must be exercised when evaluating the information provided by continuous data in the context of selection or adaptation and in determining whether they are sufficiently informative to aid in resolving relationships. Further investigation is needed to elucidate the reasons for the limitations of continuous characters, which often resolve nodes that do not exceed half the relative tree timespan. Also needed is more extensive research into what “apical” and “basal” mean in actual empirical cases.

## CONCLUSION

Continuous characters analyzed without subjective discretization by parsimony do carry phylogenetic signals to infer species relationships. However, the accuracy of phylogenetic inference depends on the underlying process of character change over time. When quantitative traits evolve under random drifting (i.e., neutral evolution), relationships are more accurately recovered at shallower and intermediate levels of cladograms, but rarely at deeper levels, regardless of the evolutionary rate applied. Other underlying evolutionary processes of character evolution (e.g., selection and adaptive radiations) generate weaker phylogenetic signals and are more useful for refining relationships of recently diverged sister species. Weighting against homoplasy has no apparent effect on cladogram inferences from quantitative traits, regardless of the strength of down-weighting. This fact suggests, among other possibilities, that quantitative traits are rarely homoplastic or that discretization minimizes the delimitation of homoplastically-distributed states. Our findings highlight that quantitative traits are sources of evolutionary information for inferring the relationships of closely related species, which can be difficult to resolve using other data sources. At lower taxonomic levels, variation is often expressed as continuous changes in morphometric sizes, proportions, and body shapes. The inclusion of quantitative traits in cladistics in such situations is therefore likely to increase the accuracy of phylogenetic hypotheses.

## DATA ACCESSIBILITY

The simulated and inferred trees, along with the character matrices analyzed in this publication, will be available for access at 10.5281/zenodo.10456412

## AUTHORS’ CONTRIBUTIONS

All authors contributed equally to the conceptualization of the study. VAT and AD assumed leadership roles in conducting data curation and formal analysis. VAT, AD, JC, and MP jointly contributed to validating the results and writing the manuscript.

## ACKNOWLEDGEMENTS

We thank our colleagues, particularly graduate students, postdoctoral researchers and faculty staff of the Museu de Zoologia da Universidade de São Paulo (MZUSP), Museu de Biodiversidade do Cerrado (MBC), and Universidade Federal de Uberlândia for their contributions with discussions that improved this manuscript.

